# RING1 dictates GSDMD-mediated inflammatory response and host susceptibility to pathogen infection

**DOI:** 10.1101/2024.11.05.622030

**Authors:** Yuanyuan Li, Wenqing Gao, Yuxin Qiu, Jiasong Pan, Qingqing Guo, Xuehe Liu, Lu Geng, Yajie Shen, Zhidong Hu, Suhua Li, Shanshan Liu, Hua Yang, Baoxue Ge, Xiaoyong Fan, Xiangjun Chen, Jixi Li

**Affiliations:** State Key Laboratory of Genetic Engineering, Department of Neurology, Huashan Hospital and School of Life Sciences, Fudan University, Shanghai, 200438, China; Centre for Immunology and Infection Control, School of Biomedical Sciences, Queensland University of Technology, Brisbane, QLD, 4059, Australia; Shanghai Institute of Infectious Diseases and Biosecurity & Shanghai Public Health Clinical Center, Fudan University, Shanghai 201508, China; Division of Natural Science, Duke Kunshan University, Jiangsu, 215316, China; Shanghai Key Laboratory of Tuberculosis, Department of Microbiology and Immunology, Shanghai Pulmonary Hospital, Tongji University School of Medicine, Shanghai 200433, China; Department of Neurology, Huashan Hospital and Institute of Neurology, Fudan University, Shanghai, 200040, China

**Keywords:** Pyroptosis, GSDMD, RING1, Ubiquitination, Bacterial infection

## Abstract

RING1 is an E3 ligase component of the polycomb repressive complex 1 (PRC1) with known roles in chromatin regulation and cellular processes such as apoptosis and autophagy. However, its involvement in inflammation and pyroptosis needs to be better characterized. Here, we found that human RING1, but not RING2, promoted K48-linked ubiquitination of Gasdermin D (GSDMD) and acted as a negative regulator of pyroptosis and bacterial infection. *Ring1* knockout mice increased the bacterial loads and mortality rate by treatment of *S. typhimurium*. Upon infection by *M. tuberculosis* (Mtb) H37Rv strain, RING1 deletion initially reduced bacterial loads but later increased lung inflammation and impaired immune defense responses in mice. Also, *Ring1* knockout exacerbated acute sepsis induced by lipopolysaccharide (LPS). Mechanistically, RING1 directly interacts with GSDMD and ubiquitinates the K51 and K168 sites of GSDMD for K48-linked proteasomal degradation, thereby inhibiting pyroptosis. Inhibition of RING1 E3 ligase activity by mutation or small molecular compound increased GSDMD level and cell death during pyroptosis. Our findings reveal that RING1 dictates GSDMD-mediated inflammatory response and host susceptibility to pathogen infection, highlighting RING1 as a potential therapeutic target for combating infectious diseases.

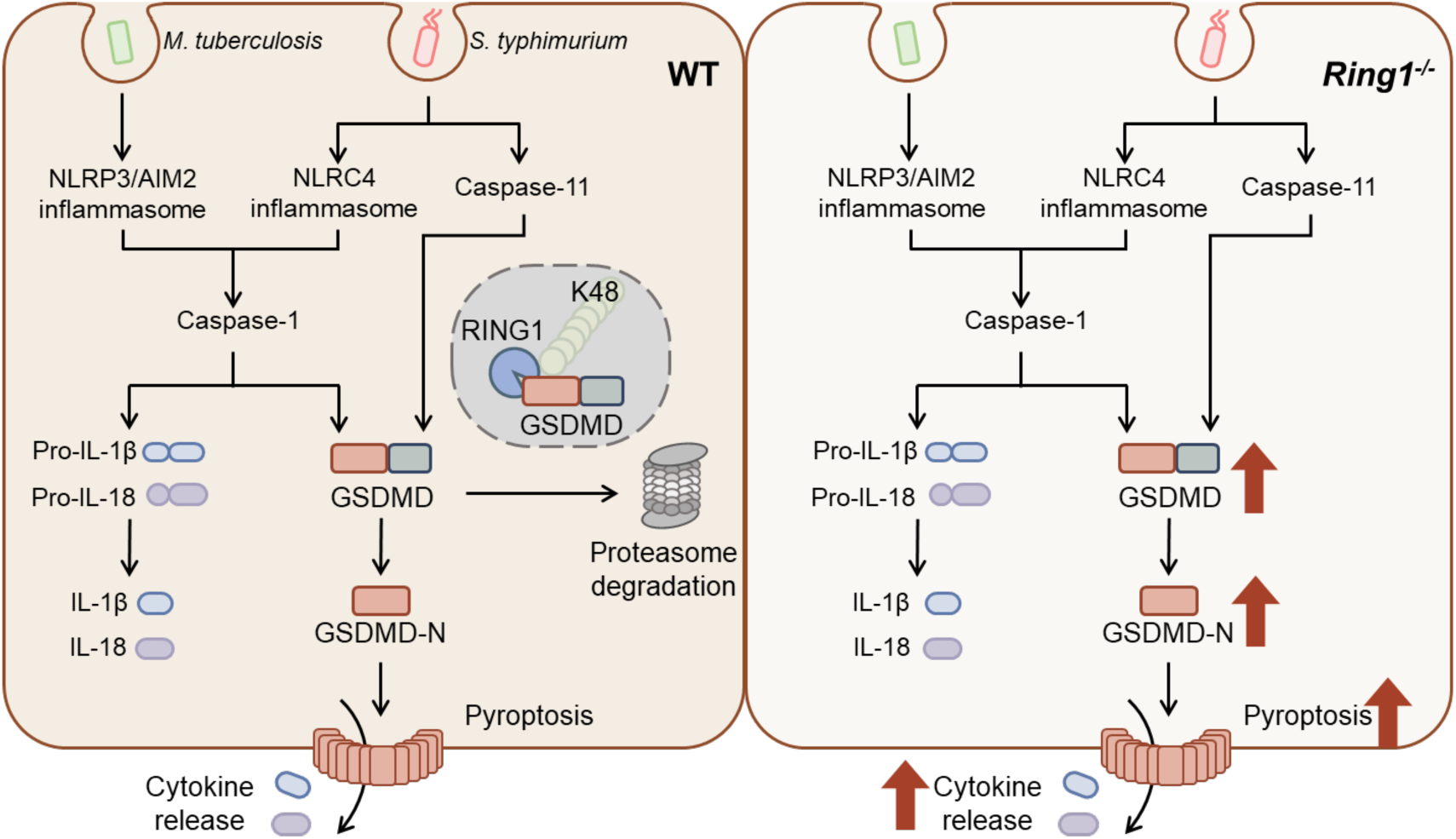
Graphic Abstract

**Highlights:**

- E3 ubiquitin ligase RING1 targets GSDMD for proteasomal degradation.
- Mice lacking RING1 are more vulnerable to *S. typhimurium* infection and LPS- induced sepsis.
- In the early stages of *M. tuberculosis* (Mtb) infection, *Ring1^-/-^* mice exhibit efficient bacterial clearance.
- Chronic inflammation in *Ring1^-/-^* mice impairs the ability to control Mtb in the late stages of infection.

## Introduction

RING1 (also known as RNF1/ RING1A) is characterized by the presence of a RING domain, and a conserved zinc-binding motif that imparts E3 ligase activity (Shen et al., 2018). RING1 is a core component of the polycomb repressive complex 1 (PRC1), which is a critical epigenetic regulator of gene expression. In detail, RING1 catalyzes the monoubiquitination of histone H2A at lysine 119 (H2AK119ub1), leading to chromatin compaction and transcriptional repression (Blackledge and Klose, 2021). Therefore, RING1 plays a crucial role in maintaining cellular identity, developmental regulation, and DNA damage response. For example, RING1 regulates neurodevelopment of the mouse telencephalon (Eto et al., 2020), and regulates DNA damage repair during neurogenesis (Ryan et al., 2024). Beyond neurogenesis, RING1 protects against colitis by regulating the mucosal immune system and maintaining colonic microbial balance (Wang et al., 2023). In blood cancers, *Ring1* blocks the gene *Glis2* to help keep AML stem cells alive in mouse models of acute myeloid leukemia (Shima et al., 2018). However, despite these advances, there is a significant gap in understanding RING1’s role in inflammation-related diseases, highlighting the need for further research to explore its potential contributions in the context of pathogen infection.

Among the various forms of programmed cell death, pyroptosis stands out as a critical pro-inflammatory mechanism in the body’s response to pathogen infection and tissue damage. Its dysregulation can contribute to various inflammatory diseases or organ damage, including sepsis (Hu et al., 2020), tuberculosis (TB) (Chai et al., 2022), *S. typhimurium* infection (Wu et al., 2022), etc. During pyroptosis, damage- or pathogen-associated molecular patterns (DAMPs/PAMPs) are recognized by various inflammasomes, encompassing both canonical and non-canonical forms. These inflammasomes then serve as molecular platforms that recruit and subsequently activate caspase-1, along with caspase-4/5 in humans and caspase-11 in mice (He et al., 2015; Kayagaki et al., 2015; Shi et al., 2015). Gasderin D (GSDMD) undergoes proteolytic cleavage upon activation by inflammatory caspases. This cleavage event generates an N-terminal fragment (GSDMD-N) responsible for membrane pore formation and a C-terminal fragment (GSDMD-C) (Kuang et al., 2017; Shi et al., 2015). Additionally, GSDMD-mediated release of cytokines, including interleukin-1β (IL-1β), and interleukin-18 (IL-18), contributes to the inflammatory response during infections and tissue damage (Xia et al., 2021).

In this study, we investigate the potential roles of RING1 in inflammatory response and host susceptibility to pathogen infection. *Ring1* knockout in mice exacerbates sepsis and acute shock induced by lipopolysaccharide (LPS) and *S. typhimurium*, respectively. Moreover, RING1 deletion enhances early-stage clearance of *M. tuberculosis* (Mtb) but aggravates its dissemination and pulmonary inflammatory response in later stages. Mechanically, RING1 targets GSDMD for proteasomal degradation, thereby acting as an inhibitor of pyroptosis. These results illuminate the intricate interplay between pyroptosis and bacterial pathogenesis, highlighting RING1 as a potential therapeutic target for combating infectious diseases.

## Results

### RING1 deletion exacerbates inflammatory response during pathogen infection

RING1 and RNF2 (RING2) are both essential components of the PRC1 and are highly homologous (55% identity). Despite this similarity, RING2 is more prominently involved in cancer-related pathways and antiviral responses, such as enhancing Wnt/β-catenin signaling, inhibiting interferon functions, and suppressing the anti-tumor activity of NK and CD4^+^ T cells (Liu et al., 2018; Peeters and DuPage, 2021; Zhang et al., 2021). Additionally, RING2 knockout models have shown embryonic lethality, highlighting its critical role in early development (Voncken et al., 2003). However, little was known about the roles of RING1 in the context of pathogen infections and inflammatory diseases. We generated *Ring1^-/-^* mice by knocking out exons 3 and 4 of the *Ring1* gene using the CRISPR/Cas9 system (**Figure S1A**). Knockout of the *Ring1* gene caused mild axial skeleton abnormalities (del Mar Lorente et al., 2000; Ryan *et al*., 2024), but the mice maintained normal body weight (**Figure S1B**). We then employed *S. typhimurium* and *M. tuberculosis* (Mtb) to establish mice models of acute and chronic inflammation, respectively. In the *Salmonella*-induced acute shock model, the absence of RING1 significantly increased the mortality rate (**Figure 1A-B**). 8 hours after i.p. injection of *Salmonella*, the bacterial load in primary tissues and cytokine levels in the serum were measured. We observed an elevation in colony-forming units (CFU) in the spleen of *Ring1^-/-^* mice, while CFU levels in the liver remained unchanged (**Figure 1C**). Moreover, IL-1β was increased in the serum of *Ring1^-/-^*mice, whereas IL-6 and IL-12p70 showed no significant changes (**Figure 1D**). Notably, IL-5 was decreased in *Ring1^-/-^* mice (**Figure 1C**), indicating that T helper cell activation was inhibited. These findings suggest that excessive inflammation from acute infection may impair the host’s ability to kill bacteria, despite severe cell death.

**Figure 1.**
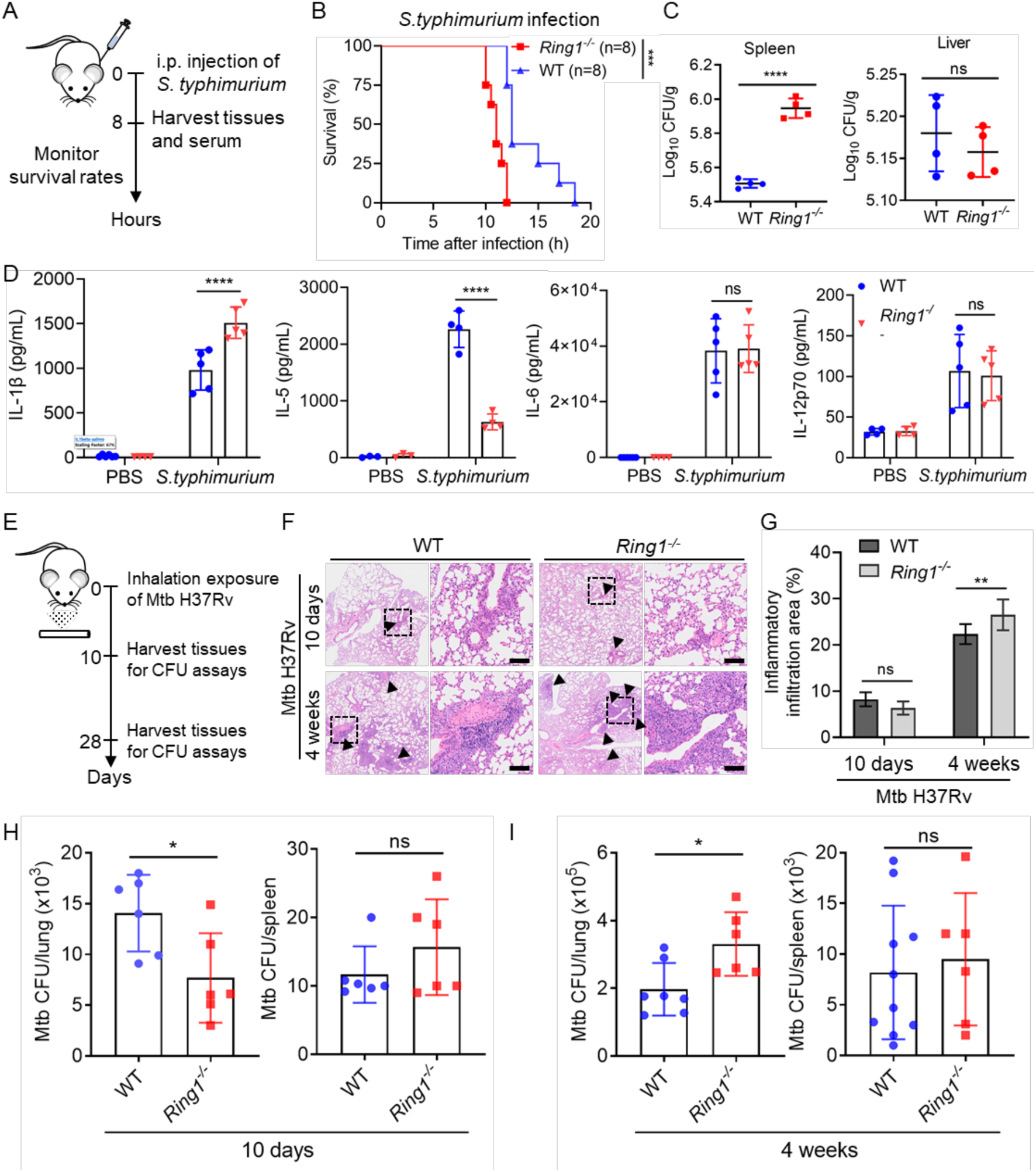
RING1 deletion increases inflammatory response upon different pathogen infection. A. The schematic diagram of experimental procedures. For *in vivo* infection, 7-week-old female mice (n=8) were i.p. injected with 5 × 10^8^ CFU of *S. typhimurium* SL14028. B. The survival rate was monitored and quantified by the Kaplan-Meier survival plot. C. The bacterial loads in the spleen and liver were analyzed 8 h after injection (n=4). D. 8 h post-infection, cytokine concentrations in the serum of WT and *Ring1^-/^*^-^ mice were measured using ELISA. E. The schematic diagram of experimental procedures. Mice were infected with Mtb H37Rv strain. F. The histopathological changes in the lungs were examined by H&E staining. Black arrows represent foci of cellular infiltration. Scale bars, 100 μm. G. Quantification of inflammatory areas in the lungs of mice treated as in F. H-I. Bacterial CFUs in lungs or spleens of mice infected with Mtb H37Rv strain. Error bars represent SD, and the statistical analysis was conducted using Two-way ANOVA (C, D and G) or Students’ *t*-test (H-I). *, *p* < 0.05; ***, *p* < 0.001; ****, *p* < 0.0001; ns, *p* > 0.05, not significant.

During Mtb infection, resident alveolar macrophages are the primary cell type that initially uptake Mtb (Cohen et al., 2018). Mtb-induced phagosome damage releases bacterial DNA and RNA into the host cytosol, activating innate immune-sensing pathways (Chai et al., 2020). Moreover, plasma membrane damage during Mtb infection triggers NLRP3 activation and pyroptosis (Beckwith et al., 2020). *Ring1^-/-^*mice exhibited increased inflammatory infiltration in the lungs at 4 weeks post-infection (**Figure 1E-G**). Interestingly, RING1 deletion initially reduced bacterial loads in the lungs at 10 days post-infection but showed a CFU elevation of Mtb at 4 weeks post-infection (**Figure 1H-I**). These findings suggest that RING1 may play a dual role in Mtb infection, initially aiding bacterial clearance but later exacerbating bacterial dissemination and lung inflammation.

### RING1 depletion aggravates LPS-induced sepsis

Given that sepsis involves acute inflammation similar to severe infections, we use LPS-induced sepsis model to explore the effects of RING1 on systemic inflammation and pyroptosis (van der Poll et al., 2021) (**Figure 2A**). Kaplan-Meier survival analysis showed a significant decrease in the survival rate of *Ring1*^-/-^ mice (**Figure 2B**). In the parallel group, euthanasia was performed 8 hours after intraperitoneal (i.p.) injection of LPS, and the levels of inflammatory factors in the serum were measured. The results revealed a significant increase of serum inflammatory cytokine levels after LPS injection (**Figure 2C-D**). The levels of IL-1β, IL-5, IL-12p70, and TNF-α in the serum of *Ring1*^-/-^ mice were higher than those of WT mice, while IL-6 and IL-10 levels were not significantly changed (**Figure 2C-D**). IL-1β and TNF-α, released mainly by activated macrophages and monocytes, play vital roles in recruiting immune cells (Shapouri-Moghaddam et al., 2018). IL-5 supports B cell and eosinophil functions, and IL-12p70 promotes T cell differentiation into Th1 cells and enhances NK cell cytotoxicity (Briukhovetska et al., 2021). Therefore, these cytokine changes suggest increased innate and adaptive immune responses in *Ring1*^-/-^ mice. To further investigate the molecular mechanisms underlying the differences in survival rates and serum cytokine levels, we performed WB analysis on spleen and lung tissues (**Figure 2E**). In the spleen, *Ring1* deletion led to an increase in cleaved caspase-1 (Cle-CASP1) and GSDMD-N (**Figure 2F**). A similar trend was observed in the lung, although the expression levels of pro-caspase-1/11 remained unchanged. Further IHC analysis revealed that LPS induced acute inflammation in the spleen, and *Ring1* knockout resulted in a significant increase in lymph node area (**Figure S1C-D**), suggesting a potential role for RING1 in modulating immune cell function and distribution. The periarterial lymphoid sheath (PALS) thickness, which is associated with the proliferation of inflammatory cells in the spleen, was significantly increased in *Ring1*^-/-^ mice in response to LPS (**Figure S1E-F**). Collectively, these observations indicate that RING1 plays a critical role in mitigating excessive inflammation and preserving tissue integrity during acute infection.

**Figure 2.**
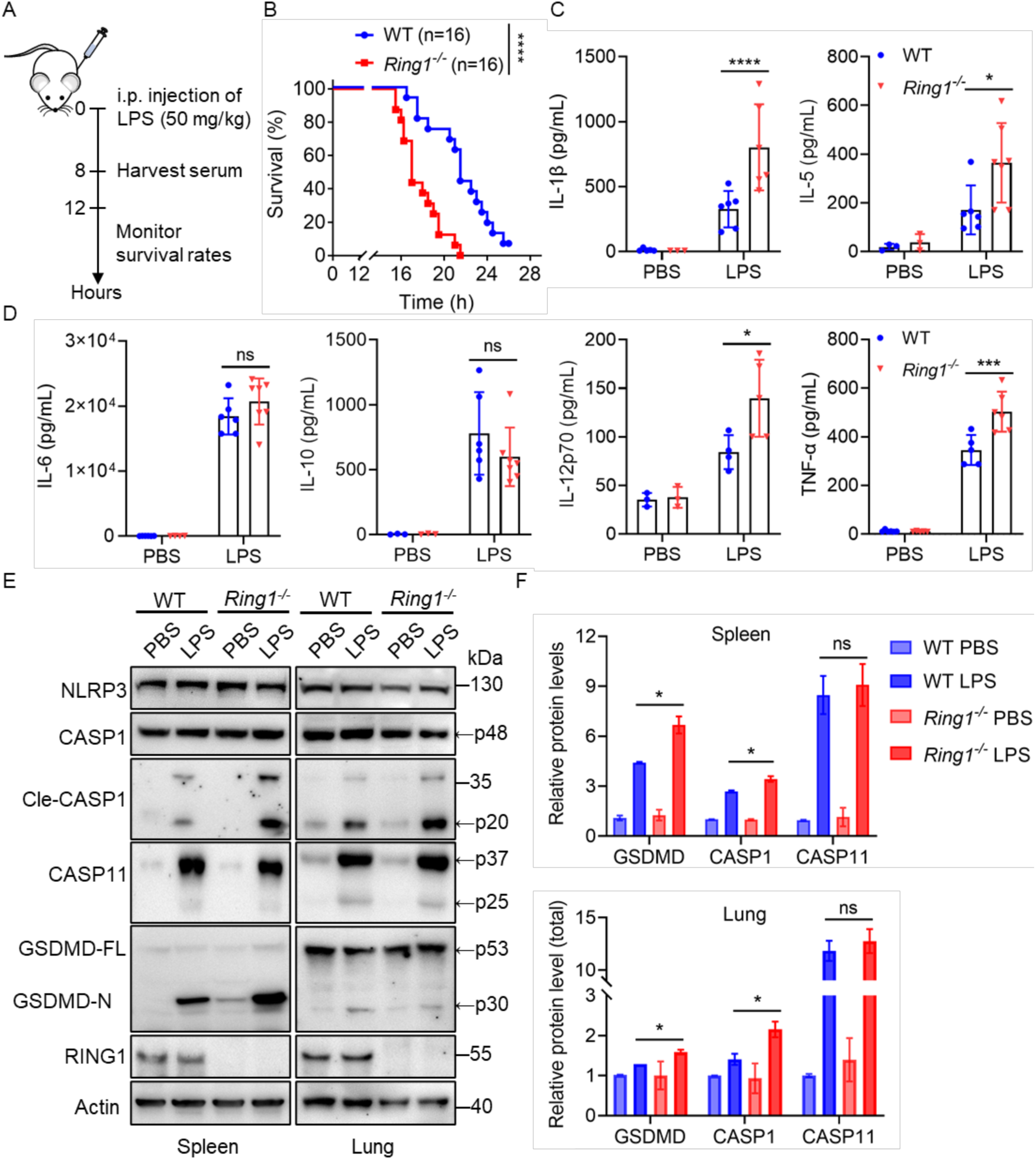
*Ring1* knockout mice are more susceptible to LPS-induced sepsis. A. The schematic diagram of experimental procedures. 6-week-old female mice (n=16) were i.p. injected with 50 mg/kg LPS or an equivalent volume of PBS. B. The survival rate was monitored and quantified by the Kaplan-Meier survival plot. C-D. At 8 h post-infection, cytokine concentrations in the serum were quantified using ELISA. E. Lung and spleen tissue lysates were analyzed via Western blotting. F. The quantification of WB results in E. Error bars represent SD, and the statistical analysis was conducted using Two-way ANOVA. *, *p* < 0.05; ***, *p* < 0.001; ****, *p* < 0.0001; ns, *p* > 0.05, not significant.

### RING1 negatively regulates GSDMD stability

To further explore how RING1 influences GSDMD and CASP1 protein, we analyzed both transcriptional and post-transcriptional regulation levels. Using reverse transcription PCR (RT-PCR), we quantitatively assessed the mRNA levels of vital pyroptosis-related genes (**Figure S2A**). The results showed no significant changes in the transcription levels of *Nlrp3*, *Caspase-1*, *Caspase-11*, or *Gsdmd* genes in *Ring1^-/-^* iBMDMs, indicating that RING1 does not regulate GSDMD and CASP1 at the transcriptional level. Next, we use cycloheximide (CHX) to inhibit protein synthesis and determine the stability of GSDMD and CASP1 proteins. When transfected with RING1-mCherry, the protein stability of CASP1 remained unchanged compared to it transfected with the empty vector control (EV) (**Figure S2B**). However, RING1 decreased the half-life of endogenous GSDMD in U251 cells (**Figure 3A-C**). Before CHX treatment, GSDMD protein levels were already down-regulated in RING1- overexpression U251 cells (**Figure 3C**). We constructed Ring1^-/-^ immortalized bone marrow-derived macrophages (iBMDMs) using the CRISPR-Cas9 system to verify this finding in immune cells. Consistent with the observation in U251 cells, GSDMD exhibited a longer half-life in *Ring1^-/-^*iBMDMs, with or without CHX treatment (**Figure 3D-F**). Moreover, the degradation of GSDMD was inhibited by MG132 treatment, which further confirms that GSDMD was degraded via the ubiquitin-proteasome system (**Figure 3G**). These results showed that RING1 functions in modulating pyroptosis by post-translational modifications of GSDMD.

**Figure 3.**
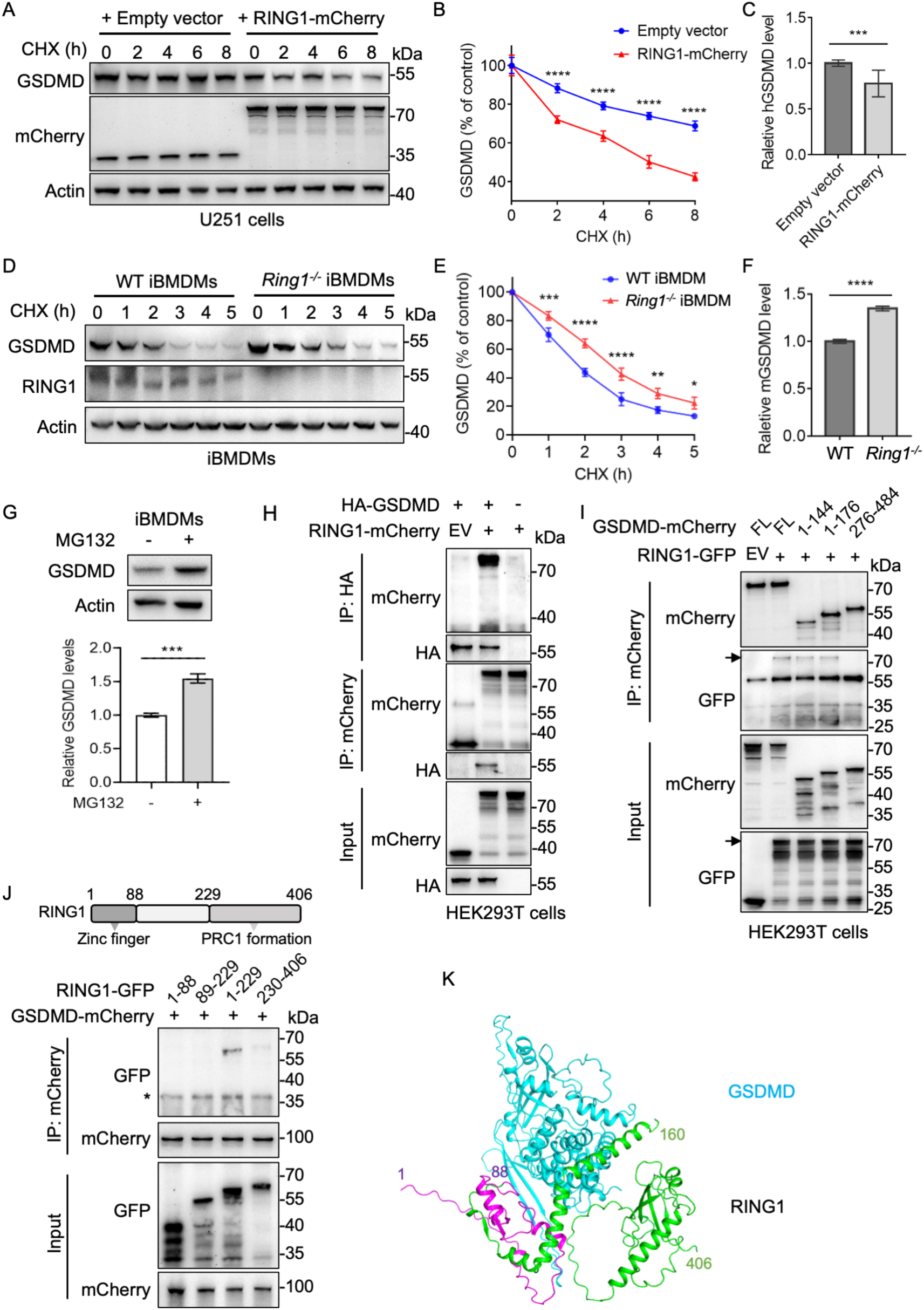
RING1 interacts with GSDMD. A. Detection of endogenous GSDMD half-life in the presence or the absence of mCherry-tagged RING1. B. GSDMD expression was quantified relative to the baseline GSDMD expression levels in unstimulated EV-transfected or RING1-mCherry-transfected cells. C. Relative amounts of GSDMD were quantified relative to the baseline GSDMD expression levels in unstimulated EV-transfected cells. D. Wild type (WT) and *Ring1* knockout iBMDMs were treated with CHX for different time intervals, followed by WB analysis. E. The quantification of WB results in panel G. F. Qualification of the relative GSDMD levels in untreated WT or *Ring1^-/-^* iBMDMs. G. iBMDMs were treated with 10 μM MG132 for 4 h, and then subjected to WB. The relative GSDMD protein levels were quantified. H. Co-IP analysis of HA-GSDMD and RING1-mCherry. I. GSDMD-mCherry or its mutants and RIGN1-GFP were co-transfected into HEK293T cells, followed by WB analysis. Arrows point to the RING1-GFP bands, and the asterisk indicates the IgG heavy chain. J. Co-IP experiment of RIGN1-GFP or its mutants and GSDMD-mCherry. The asterisk indicates the IgG light chain. K. The predicted RING1-GSDMD complex structure using the AlphaFold3 server (https://www.alphafoldserver.com/) was shown with a cartoon model. Human GSDMD is shown in cyan, and RING1 is shown in purple (residues 1-88) and green (residues 89-160 and 258-406) colors. The flexible region (residues 161-257) was invisible. Error bars represent standard deviation (SD). The statistical analysis was performed using Two-way ANOVA (E and H), or Students’ *t*-test (F and I). *, *p* < 0.05; **, *p* < 0.01; ***, *p* < 0.001; ****, *p* < 0.0001.

### E3 ligase RING1 directly interacts with GSDMD

Based on these findings, we performed the Co-IP experiment to validate the physical interaction between GSDMD and RING1 (**Figure 3H-J**). When HA-tagged GSDMD was co-transfected with mCherry-tagged RING1, GSDMD co-immunoprecipitated with RING1 but not with mCherry empty vector (EV) (**Figure 3H**). To avoid the cytotoxic effects of GSDMD-N (1-276), we constructed truncated versions of GSDMD-N. RING1 specifically interacted with the full-length GSDMD and the N-terminus (residues 1-144 or 1-176) but not with the C-terminus (**Figure 3I**), implying that the conformation change of GSDMD-N after cleavage by inflammatory caspases is unnecessary for binding with RING1 (Devant and Kagan, 2023). Like other RING-type E3 ligases, RING1 contains a Zinc-finger domain (resiudes 48-88) and a region necessary for interaction with CBX2 (residues 230-406). To map the crucial domains responsible for the interaction with GSDMD, we constructed a series of GFP-tagged RING1 truncated mutants. GSDMD specifically interacted with RING1 (1-229) truncation, but not the 1-88 or the 89-229 domain, whereas GSDMD interacted weakly with the C-terminal domain of RING1 (230-406) (**Figure 3J**), indicating that the stereochemical conformation of RING1 (1-229) is needed for binding with GSDMD. Next, the AlphaFold3 server was used to predict the possible interaction surfaces between GSDMD and RING1. The predicted complex structure showed that GSDMD interacted tightly with the N-terminal domain of RING1, similar to the above result (**Figure 3K**). To further investigate whether endogenous GSDMD colocalizes with RING1, HeLa and U251 cells were subjected to immunofluorescence analysis (**Figure S2C**). The quantification of fluorescence density indicated that GSDMD colocalized with RING1, suggesting that RING1 might play a vital role in GSDMD regulation (**Figure S2D**). Together, the above data indicated that GSDMD directly interacts with RING1.

### RING1 catalyzes K48-linked polyubiquitination of GSDMD

As RING1 might affect the stability of GSDMD protein via the proteosome pathway (**Figure 3A-G**), iBMDM cells were induced to undergo pyroptosis with LPS plus nigericin (Nig) for investigating the ubiquitination status of GSDMD. Detection of the immunoblotting for Ub after immunoprecipitating GSDMD showed that ubiquitination occurred in early states of pyroptosis (**Figure S2E**). To further investigate how RING1 mediates the post-translational regulation of GSDMD, we explored its specific role in catalyzing ubiquitination. GSDMD and His-Ub (WT, K48 only, K63 only) were transfected into HEK293T cells with the presence or absence of RING1. The results showed that RING1 mainly catalyzed the formation of K48-linked polyubiquitin chains onto GSDMD (**Figure 4A**). During ATP-induced pyroptosis, RING1 deletion also reduces the K48-linked ubiquitination of GSDMD in iBMDMs (**Figure 4B**). Next, to demonstrate whether RING1 could synthesize Ub chains directly to GSDMD, an *in vitro* ubiquitination assay was performed by adding the purified proteins human RING1, GSDMD, E1 enzyme UBA1, E2 enzyme UBE2D3, and ubiquitin. A ladder of high-molecular-weight bands on GSDMD indicated that RING1 ubiquitinated GSDMD only when RING1 was present (**Figure 4C**).

**Figure 4.**
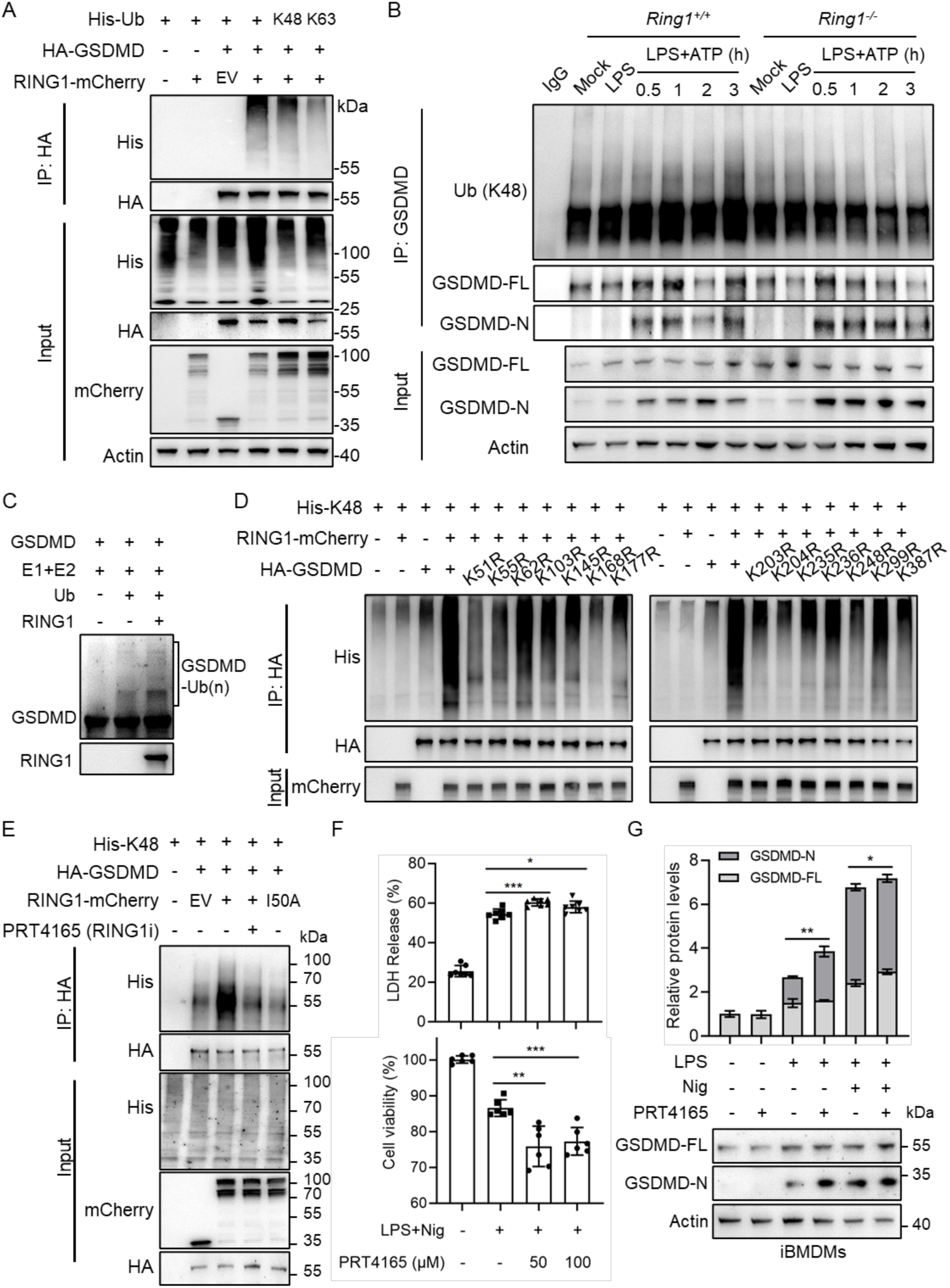
GSDMD is ubiquitinated by RING1. A. His-tagged Ub wild-type or its mutants (K63-only and K48-only) were transfected into HEK293T cells with indicated plasmids. Then, cells were subjected to HA pull-down and IB analysis. B. WT or *Ring1^-/-^* iBMDMs were treated with LPS and ATP. Cell lysates were immunoprecipitated with anti-GSDMD antibody and then subjected to IB analysis. C. *In vitro* ubiquitination assay for GSDMD. Purified GSDMD was incubated with indicated proteins. D. RING1-mCherry, His-K48, and HA-GSDMD or its mutants were co-transfected into HEK293T cells. Cell lysates were immunoprecipitated with anti-HA antibody and then analyzed with WB. E. HEK293T cells were transfected with indicated plasmids. 16 h after transfection, cells were treated with 50 μM PRT4165 for 1 h. F. iBMDMs were treated with the indicated drugs and analyzed by Western blot. Total GSDMD protein levels are quantified in the upper panel. G. iBMDMs were treated with LPS and Nig for 2 h, with or without PRT4165, and then analyzed for cell viability and LDH release. Error bars represent SD, and the statistical analysis was conducted using the Students’ *t*-test. *, *p* < 0.05; **, *p* < 0.01; ***, *p* < 0.001; ****, *p* < 0.0001.

To further investigate the potential ubiquitination sites of RING1, all 15 lysine residues of GSDMD were mutated to arginine individually. Since lysine (K) and arginine (R) are basic amino acids, mutating lysine residues to arginine residues can inhibit lysine-mediated ubiquitination while retaining the charge properties. However, the K43R, K103R, and K168R mutant expression levels were significantly reduced (**Figure S3A**). This decrease in expression may be due to mRNA destabilization or disruption of critical protein structures needed for stability and function. The same amount of IP-enriched GSDMD was loaded for each sample, and the ubiquitination levels of GSDMD were detected by WB. As depicted in **Figure 4D**, K48-linked ubiquitination of GSDMD was attenuated at multiple mutation sites, with the most notable effects observed in the mutants at K168 and K204. In addition, the ubiquitination of GSDMD was increased by WT RING1 but not by catalytic inactive RING1 or under treatment by RING1 inhibitor PRT4165 (**Figure 4E**). This result indicated that RING1 ubiquitinates GSDMD in an E3 ligase activity-dependent manner.

### RING1 deficiency promotes pyroptosis in macrophages

To investigate the potential roles of RING1, we induced NLRP3 inflammasome-mediated pyroptosis in iBMDMs using LPS and Nig. Inhibition of RING1 E3 ligase activity increased the cell death rate of iBMDMs (**Figure 4F**). Also, treatment of PRT4165, a BMI1/RING1 inhibitor, elevated the GSDMD protein level during pyroptosis (**Figure 4G**). These findings suggest that RING1’s regulation of pyroptosis relied on its E3 ligase activity. Next, we assessed cell death rates and the expression of pyroptosis-related proteins in *Ring1*^-/-^ iBMDMs upon PRT4165 treatment. Compared to wild-type iBMDMs, *Ring1* knockout cells exhibited increased sensitivity to pyroptotic stimuli, with a significant increase in cell death rate and a decrease in cell survival rate (**Figure 5A**). Validation of time-course drug treatment showed that the survival rate of *Ring1*^-/-^ iBMDMs significantly decreased (**Figure 5B**). Meanwhile, real-time confocal microscopy imaging showed that the morphology of pyroptosis in *Ring1*^-/-^ iBMDMs is not different from that of wild-type cells (**Figure S3B**). Next, primary bone marrow-derived macrophages (BMDMs) from wild-type and knockout mice were obtained. PI staining showed that *Ring1*^-/-^ BMDMs exhibited a significant increase in cell death rate during ATP-induced pyroptosis (**Figure 5C and Figure S3C**). Furthermore, either CY-09 (an NLRP3 inhibitor) or VX765 (a CASP1 inhibitor) alleviated the increased cell death of *Ring1*^-/-^ iBMDMs, but only the GSDMD inhibitor disulfiram (DSF) could restore the cell viability rate to the level of WT iBMDMs (**Figure S3D**). This result indicates that the effect of *Ring1* deletion on pyroptosis is primarily dependent on GSDMD.

**Figure 5.**
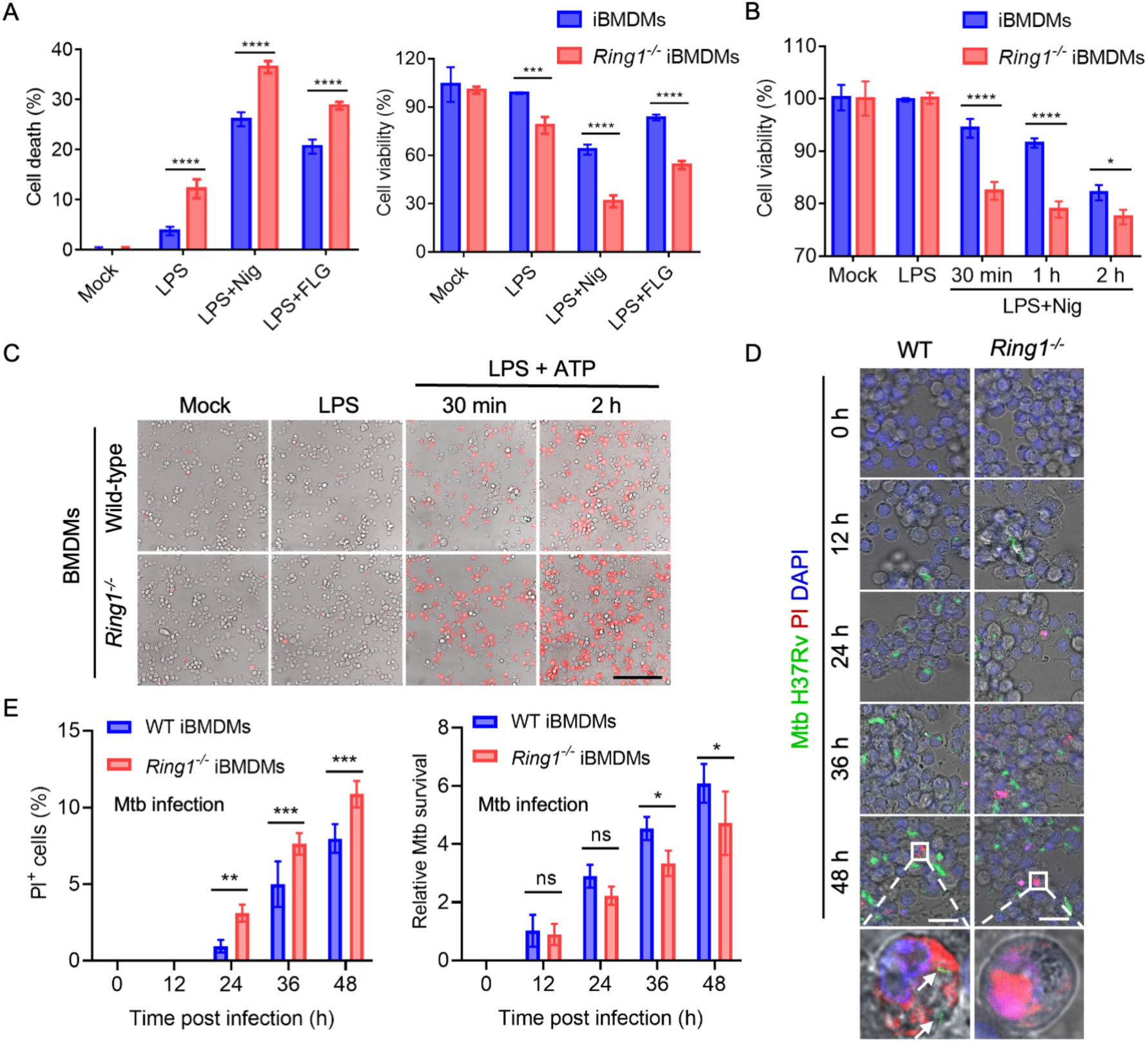
*Ring1* knockout macrophages are more sensitive to pyroptosis. A-B. WT and *Ring1*^-/-^ iBMDMs were pre-treated with 2 μg/mL LPS for 2 hours, followed by continued treatment with 40 μM Nigericin (Nig) or 50 μM Flagellin (FLG) for 2 hours or the indicated time. The cell death rate was calculated by assessing LDH release, and cell viability was determined by CCK-8 assay. C. WT and *Ring1*^-/-^ primary BMDMs were pre-treated with 1 μg/mL LPS for 2 hours, followed by 5 mM ATP treatment. After drug treatment, cells were stained with propidium iodide (PI) and subjected to confocal microscopy imaging. Each dataset was generated from counting at least five random areas. Scale bar, 50 μm. D. Confocal microscopy imaging of WT or *Ring1^-/-^* iBMDMs, which were infected with GFP-expressing Mtb H37Rv and then cultured for the indicated times. The arrows mark Mtb in pyroptotic cells. Scale bars, 25 μm. E. Qualification of PI^+^ cells and relative Mtb survival treated as in D. Statistical analysis was performed using Two-way ANOVA. *, *p* < 0.05; **, *p* < 0.01; ***, *p* < 0.001; ****, *p* < 0.0001; ns, not significant.

To further investigate the role of RING1 in bacterial infection, we used Mtb H37Rv strain to infect macrophages and performed confocal imaging. Some Mtb bacilli were killed immediately after pyroptosis, while the rest were trapped in the pyroptotic cell (**Figure 5D**). The bacteria trapped in the ghost cells may be phagocytosed by neighboring cells, contributing to the local spread of infection (Beckwith *et al*., 2020). Next, we quantified fluorescence to measure the cell death rate marked by PI^+^ and the bacterial density represented by GFP green fluorescence. The results showed that the *Ring1*^-/-^ iBMDMs exhibited a higher death rate 24-48 hours after Mtb infection (**Figure 5E**). Correspondingly, the bacterial survival rate also decreased (**Figure 5E**). This may be due to the higher levels of GSDMD-N, enhancing the bacterial clearance in an early stage of infection, which is consistent with the decreased Mtb CFU shown in **Figure 1F**. Taken together, these findings indicate that RING1 is a negative regulator of GSDMD, and *Ring1* knockout promotes pyroptosis mediated by multiple inflammasomes.

### *Ring1* knockout elevates the protein expression level of GSDMD

To further reveal the effect of RING1 deficiency on cell pyroptosis, we examined the protein expression levels involved in the pyroptotic pathway at different time points (**Figure 6A**). Following treatment with Nig, the NLRP3 protein increased in both wild-type and knockout cells. With prolonged exposure to the drug, the expression of RING1 protein also gradually increased, suggesting the involvement of RING1 in the process of cell pyroptosis (**Figure 6A**). LPS treatment induces the initiation step of the NLRP3 inflammasome-mediated canonical inflammatory response pathway (Huang et al., 2021). Consistently, LPS pre-treatment increased the protein level of procaspase-1. More importantly, both in the resting state and upon stimulation, RING1 deficiency significantly increased the total protein levels of GSDMD (**Figure 6B**). ELISA of cell culture supernatants showed that *Ring1*^-/-^ iBMDMs released more IL-1β, interleukin 18 (IL-18), and TNF-α during pyroptosis (**Figure 6C-D**). However, IL-6 level was decreased in *Ring1*^-/-^ iBMDMs (**Figure 6D**), which contradicts the typical inflammatory response (Taru et al., 2024). To explore the cause of this IL-6 reduction, we conducted RNA sequencing (RNA-seq) analysis of *Ring1*^-/-^ and wild-type iBMDMs with three different treatments. Differentially expressed genes (DEGs) between *Ring1*^-/-^ and wild-type cells were identified under each condition (**Figure S4A-C**), and a GO analysis was performed on the intersecting DEGs across the three conditions (**Figure S4D-E**). The analysis revealed that genes primarily associated with RING1 were related to forebrain development and positive regulation of epithelial cell migration. These genes were also linked to cellular components such as sarcolemma, membrane regions, membrane microdomains, and membrane rafts. Additionally, IL-6 mRNA was downregulated, indicating transcriptional inhibition (**Figure S4D**). This reduction was also confirmed by the qPCR results (**Figure 3A**), which explains the decrease in IL-6 levels despite the heightened pyroptotic response observed in *Ring1*^-/-^ iBMDMs. Together, the above results indicated that *Ring1* deletion promotes pyroptosis by upregulating the GSDMD protein level.

**Figure 6.**
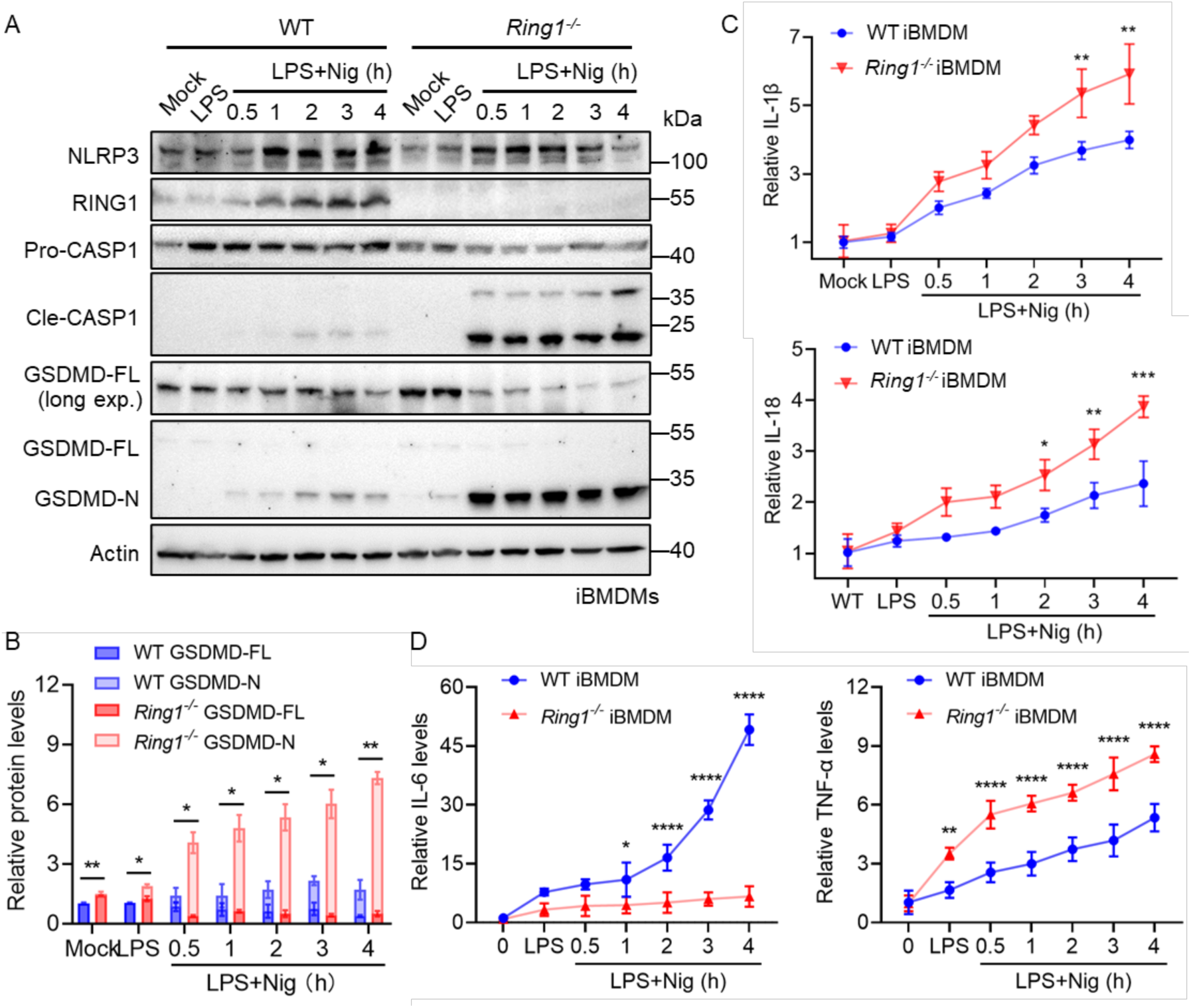
*Ring1* knockout upregulates GSDMD protein levels. A. WT and *Ring1*^-/-^ iBMDMs were pre-treated with 1 μg/mL LPS for 2 hours, followed by 20 μM Nig for various duration. Cell lysates were further subjected to WB analysis. B. Quantification of total GSDMD protein levels was performed. C-D. The cytokines (IL-1β, IL-6, IL-18, and TNF-α) released into the culture media were analyzed using ELISA. Error bars represent SD. The statistical analysis was conducted using Two-way ANOVA. *, *p* < 0.05; **, *p* < 0.01; ***, *p* < 0.001; ****, *p* < 0.0001.

## Discussion

RING1 has emerged as a critical regulator of diverse cellular processes beyond transcriptional control, encompassing the development of nervous systems, apoptosis, tumorigenesis and tumor development (Pierce et al., 2018; Zhu et al., 2020). Additionally, RING1 catalyzes the proteasomal degradation of tumor protein (P53) through K48-linked ubiquitination, thereby influencing apoptosis and senescence (Shen *et al*., 2018). Our current discovery unveils the critical roles of RING1 in inflammatory diseases.

In *S. typhimurium* infection, pyroptosis is triggered by the activation of the NLRC4 inflammasome, promoting bacterial clearance by inducing inflammatory cell death in macrophages, neutrophils, and other immune cells, thereby limiting bacterial replication and spread (Wu *et al*., 2022). RING1 deletion increased the acute inflammatory responses in mice but reduced their ability to clear bacteria (**Figure 1A- D**). Upon Mtb infection, the bacteria are initially engulfed by macrophages, leading to either latent or active TB (Chandra et al., 2022). Therefore, early clearance of Mtb is crucial for infection control. Our findings reveal that the absence of *Ring1* initially enhances the clearance of Mtb, suggesting a potential therapeutic benefit (**Figure 1E- I**). However, in later stages of infection, the subsequent pulmonary inflammation associated with *Ring1* deficiency may promote Mtb migration or proliferation (**Figure 1I**). These findings suggest that inhibiting pyroptosis during acute infections like sepsis could help reduce tissue damage. In contrast, activating pyroptosis in chronic infections may aid in bacterial clearance.

Ubiquitination regulates cellular processes by covalently attaching ubiquitin molecules to target proteins (Liu et al., 2005). This modification serves as a molecular switch, effectively regulating their stability, subcellular localization, and functional activities (Mooney and Sahingur, 2021). K48-linked ubiquitination is renowned for its canonical role in marking proteins for proteasomal degradation, influencing multiple cell death pathways (Luchetti et al., 2021; Mei et al., 2021; Shen *et al*., 2018). The role of ubiquitination in the regulation of GSDMD has emerged as a crucial regulatory mechanism that modulates its activity and function. Several E3 ubiquitin ligases have been implicated in mediating GSDMD ubiquitination (Gao et al., 2022; Luchetti *et al*., 2021; Shi et al., 2022). SYVN1 and TRIM21 can mediate human GSDMD trafficking via the K27- or K33-linked ubiquitination, respectively, while bacterial IpaH7.8 can degrade human GSDMD in the K48-linked ubiquitination (Gao *et al*., 2022; Luchetti *et al*., 2021; Shi *et al*., 2022). The key finding of this study is the identification of the first human ubiquitin ligase that degrades GSDMD. RING1 binds GSDMD and inhibits GSDMD stabilization, catalyzing K48-linked ubiquitination of GSDMD, thus inhibiting NLRP3/NLRC4-mediated pyroptosis in macrophages (**Figure 3**).

After treatment with Nig, the levels of cleaved Caspase-1 (Cle-CASP1) increased significantly in *Ring1^-/-^* iBMDMs, and there was a similar trend in the spleens of septic mice (**Figure 2E and Figure 6A**). However, RING1 does not affect CASP1 stability or gene transcription. This suggests that more GSDMD-N in *Ring1^-/-^*cells may enhance CASP1 activation through positive feedback, or *Ring1* knockout could influence the activity of upstream regulators, amplifying the Cle-CASP1 level. We also analyzed the genes associated with *Ring1* knockout using the RNA-seq method (**Figure S4**). Given RING1 functions in development and cell migration, these genes may also influence inflammatory responses. For instance, the regulation of epithelial cell migration is crucial during inflammation, as it facilitates tissue repair and barrier restoration (Nowarski et al., 2017). Additionally, the association of these genes with membrane microdomains and rafts suggests their involvement in cell polarity, cell death, and inflammation (Balakrishnan and Kenworthy, 2024; Hong et al., 2024). Therefore, it is likely that the functions of these RING1-related genes extend to modulating inflammatory responses. In conclusion, our study elucidated that RING1 dictates GSDMD-mediated inflammatory response and host susceptibility to different pathogen infection. Thus, RING1 may serve as a potential therapeutic target for the management of inflammatory diseases.

## Materials and Methods

### Mice

All animal experiments were performed in accordance with the NIH Guide for the Care and Use of Laboratory Animals, with the approval of the Scientific Investigation Board of School of Life Sciences, Fudan University (2020-JS-016). C57BL/6J mice were obtained from the SLAC Laboratory Animal Co. (Shanghai, China). Mutant alleles were backcrossed with C57BL/6J mice for more than six generations. All mice utilized in the experiments were aged between 6 to 8 weeks and weighed between 20 to 25 g.

To create the *Ring1* knockout mice, the CRISPR/Cas9 system was employed. Specifically, guide RNAs (gRNAs) were designed to target the third and fourth exons of *Ring1* gene. The sequences of the gRNAs were as follows: gRNA1: 5’- TTCCTGGCAGGCCTCTAAGCAGG-3’, gRNA2: 5’-ACCCTTGGCTTATATGTTGCTGG-3’. Fertilized mouse embryos were microinjected with the CRISPR/Cas9 system along with the synthesized gRNAs. Following microinjection, these embryos were implanted into pseudopregnant female mice. The resultant pups were subjected to screening for gene editing events and further breeding.

### LPS-induced sepsis

Female mice at 6-8 weeks of age were injected intraperitoneally with 50 mg/kg *E. coli* O111:B4 LPS (Sigma, USA) to induce sepsis. For the survival study, mice were constantly monitored for 26 hours, and then euthanized by cervical dislocation. In a separate experiment, mice were sacrificed on 8 h after LPS injection, followed by an ELISA assay from serum samples and a western blot analysis of kidney and lung tissues. Serum was prepared from coagulated blood centrifuged at 2,000 × g for 90 s at room temperature for cytokine analysis.

### Cell culture and treatment

HeLa, HEK293T, U251, iBMDM, and BMDM were cultured in high-glucose DMEM supplemented with 10% (v/v) FBS (Hyclone, UK). HeLa and HEK 293T cell lines were purchased from ATCC (Manassas, USA), and U251 cells were purchased from Mingzhoubio (Zhejiang, China). iBMDMs were kindly provided by Dr. F. Shao (NIBS, China). All cells were grown at 37°C in a 5% CO_2_ incubator (Thermo Fisher, USA). PolyJet (Signage, USA) or Lipofectamine RNAiMAX (Invitrogen, USA) was used for the transfection of corresponding LPS, plasmids, or siRNAs into cells. To induce pyroptosis, cells were pre-treated with 1 μg/mL lipopolysaccharide (LPS) for 2 hours. Subsequently, cells were treated with either 20 μM Nigericin (Nig) or 5 mM adenosine triphosphate (ATP) for indicated durations to activate the NLRP3 inflammasome. Additionally, cells were also treated with 30 μM bacterial flagellin (FLG) to activate the NAIP/NLRC4 inflammasome.

### Plasmids

Human RING1 cDNA was a kind gift from Prof. Jiahuai Han (Xiamen University, China) and cloned into the pEGFP-N1 and pmCherry-N1 vectors, respectively. HA-tagged GSDMD and mCherry-tagged GSDMD were subcloned from pcDNA3.1- GSDMD-Flag as described previously (Gao *et al*., 2022). Point mutants (K43R, K51R, K55R, K62R, K103R, K145R, K168R, K177R, K203R, K204R, K235R, K236R, K248R, K299R and K387R of GSDMD; I50A of RING1) were generated using the KOD-Plus-Mutagenesis kit (Toyobo, Osaka, Japan). The plasmids for His-tagged WT ubiquitin or its mutants (K63, K48, K33, K29, K27, K11, and K6) were kindly provided by Dr. Ping Wang (Tongji University, China). Truncated mutants of RING1 or GSDMD were amplified from the full-length RING1 and GSDMD, which were then subcloned into the pEGFP-N1 or pmCherry-N1 vectors. All constructs were confirmed by DNA sequencing.

### RT-PCR analysis

Total RNA was obtained from cells using TRIzol (Invitrogen) according with the manufacturer’s instructions. cDNA was synthesized from total RNA using PrimeScript 1st strand cDNA Synthesis Kit (6110A, TaKaRa) and analyzed by PCR using SPRY Green PCR Kit (MedChemExpress) and the following primers: mRING1-Forward: 5′-AGAATGCCAGCAAAACGTGG-3′, mRING1-Reverse: 5′-AGATAGGGCACATGAGTTCTGA-3′; GAPDH-Forward: 5′-CATGAGAAGTATGACAACAGCCT-3′, GAPDH-Reverse: 5′-AGTCCTTCCACGATACCAAAGT-3′; mGSDMD-Forward: 5′-GGTGCTTGACTCTGGAGAACTG-3′, mGSDMD-Reverse: 5′-GCTGCTTTGACAGCACCGTTGT-3′; mCaspase1-Forward: 5′-GGCACATTTCCAGGACTGACTG-3′, mCaspase1-Reverse: 5′-GCAAGACGTGTACGAGTGGTTG-3′; mCaspase11-Forward: 5′-GTGGTGAAAGAGGAGCTTACAGC-3′, mCaspase11-Reverse: 5′-GCACCAGGAATGTGCTGTCTGA-3′; mNLRP3-Forward: 5′-TCACAACTCGCCCAAGGAGGAA-3′, mNLRP3-Reverse: 5′-AAGAGACCACGGCAGAAGCTAG-3′. mIL6-Forward: 5′-TACCACTTCACAAGTCGGAGGC-3′, mIL6-Reverse: 5′-CTGCAAGTGCATCATCGTTGTTC-3′.

### Bacterial pathogen infection

*S. typhimurium* (SL14028s) was a kind gift from Prof. Yufeng Yao (Shanghai Jiao Tong University) and was cultivated in LB media. For injection, log-phase bacteria were centrifuged, washed, and resuspended in phosphate-buffered saline (PBS) to achieve a concentration of 1×10^9^ cells/mL. Mice were intraperitoneally (i.p.) injected with 5×10^8^ cells per mouse.

Mtb H37Rv, which expresses EGFP (Mtb-EGFP) continuously, was cultured in Middlebrook 7H9 medium supplemented with 0.05% Tween-80 and 10% OADC (BD Biosciences) until the bacterial optical density (OD_600_) reached approximately 0.6. Macrophages were seeded in six-well plates at a density of 5×10^5^ cells per well and precultured in DMEM medium supplemented with 10% FBS for 12 hours before infection. Mtb was washed twice using PBS containing 0.05% Tween-80, followed by centrifugation to pellet the bacteria. The pellet was resuspended thoroughly in DMEM medium with 0.05% Tween-80. For mouse infections, Mtb in the logarithmic phase was resuspended in PBS to an OD_600_ of 0.1. A 5 mL portion of this suspension was loaded into the aerosol system for inhalation exposure, allowing the infection of six-week-old SPF C57BL/6J mice through the aerosol route. At 10 days and 4 weeks post-infection, randomly selected mice were euthanized, and their lung and spleen tissues were collected for colony-forming unit (CFU) determination and histological analysis using HE staining.

### Measurement of cell death and viability

To measure the percent of cell death, the CytoTox 96 Non-Radioactive Cytotoxicity Assay Kit (Promega) was used to quantify the release of lactate dehydrogenase (LDH). Cell viability was determined by the Cell Counting Kit-8 (Beyotime). All the specific steps were carried out according to the manufacturer’s instructions. Luminescence and absorbance were measured on a SpectraMax M5 plate reader.

### Confocal analysis

Images were captured using a Leica TCS SP8 DIVE FALCON laser scanning confocal microscope and analyzed using a Leica Application Suite X software. Density was calculated using ImageJ. All images are representative of at least three independent experiments.

### Protein preparation, immunoprecipitation, and immunoblotting analysis

For ubiquitination analysis, the total cellular protein was isolated with a lysis buffer containing 50 mM Tris-HCl, pH 7.5, 150 mM NaCl, 1 mM EDTA, 2% SDS, and complete protease inhibitor cocktail tablets (Roche, Switzerland). For IP, whole-cell extracts were lysed in a buffer composed of 50 mM Tris-HCl (pH 7.5), 150 mM NaCl, 1 mM EDTA, and 1% NP-40 containing protease inhibitors. After prewashing with corresponding IgG (Santa Cruz Biotechnology) and protein A/G agarose (Beyotime Inst Biotech, China) for 1 h at 4 °C, cell lysates were incubated with the appropriate primary antibody overnight at 4 °C followed by 4 h incubation with protein A/G agarose beads. The beads were washed five times with the lysis buffer and were eluted in a 5×SDS/PAGE loading buffer for immunoblotting. For immunoblotting analysis, cells were lysed with a buffer (50 mM Tris-HCl, pH 7.5, 150 mM NaCl, 1 mM EDTA and 1% Triton X-100) supplemented with protein inhibitors. The cell lysates were incubated with the following antibodies: anti-HA (Proteintech, 51064-2- AP), anti-FLAG (Proteintech, 20543-1-AP), anti-His (Proteintech, 66005-1-Ig), anti-GFP (Proteintech, 50430-2-AP), anti-mCherry (Proteintech, 26765-1-Ig), anti-human GSDMD (Abcam, ab210070; CST 93709), anti-GSDMD-N (Abcam, ab215203), anti-mouse GSDMD (Abcam, ab209845), and anti-RING1 (CST, 2820). Western blotting was performed in three independent experiments.

### Bulk RNA-seq analysis

WT and *Ring1^-/-^* iBMDM cells were treated under three conditions: Mock (solvent control), LPS (1 µg/mL for 2 hours), and LPS followed by Nigericin (20 µM for an additional 2 hours). Total RNA was using the TRIzol reagent (Invitrogen) according to the manufacturer’s protocol. Libraries were prepared according to the manufacturer’s protocol for Illumina NovaSeq 6000 platforms, and sequencing was performed on the HiSeq platform. Differential gene expression analysis was conducted between WT and *Ring1^-/-^* cells for each treatment condition. Genes with a log2 fold change (Log_2_FC) > 1 or < −1 and a *p*-value < 0.05 were considered significantly differentially expressed. Volcano plots, heatmaps, and pathway enrichment analyses were generated using the R packages ggplot2 and heatmap to visualize the differentially expressed genes (DEGs).

### Recombinant protein purification and in vitro ubiquitination assay

The full-length GSDMD, RING1, ubiquitin, E1 (UBA1), and E2 (UBE2D3) were subcloned into the pET28a vector. All proteins were expressed in *E.coli* BL21 (DE3) cells and further purified by Ni-affinity chromatography. For *in vitro* ubiquitination assay, 10 μg of purified GSDMD was mixed in a total volume of 50 μL containing 5 mM ATP, 5 mM MgCl_2_, 10 μg ubiquitin, 2 μg E1, and E2 enzymes in the presence or absence of 5 μg RING1. After incubation at 37°C for 1 h, reactions were stopped by adding 5× loading buffer and analyzed by western blotting using anti-GSDMD and anti-RING1 antibodies.

### Statistical analysis

All experiment data were analyzed using GraphPad Prism 8.0 (GraphPad software Inc. USA) and were presented as the mean ± SD. Statistical analysis was performed using Student’s t-test, one-way ANOVA or two-way ANOVA. A value of P < 0.05 was considered statistically significant.

## Acknowledgments

This work was supported by grants from the National Natural Science Foundation of China (82071782, 31670878, 32161160323, 2018M641921), the National Key Research and Development Project of China (2021YFC2301500), and the Shanghai Committee of Science and Technology (20XD1400800, 22YF1403400).

## Author Contribution

J.L. conceived and designed the study. Y.L., W.G., Y.Q., J.P., Q.G., X.L., L.G., Y.S., Z.H., S.L., S.L., H.Y., B.G., X.F., and X.C. performed the experiments and analyzed the data. Y.L. and J.L. analyzed the data and wrote the manuscript. All authors discussed the results and commented on the manuscript.

## Ethics Statement

All animal experiments were performed in accordance with the NIH Guide for the Care and Use of Laboratory Animals, with the approval of the Scientific Investigation Board of School of Life Sciences, Fudan University (2020-JS-016).

## Data and materials availability

The RNA-seq data generated in this study have been deposited in the NCBI Sequence Read Archive (SRA). Raw data including fastq files have been uploaded under the accession numbers PRJNA1165469, PRJNA1165471, PRJNA1165473, PRJNA1165474, PRJNA1165475, PRJNA1165477, PRJNA1165479, and PRJNA1165480. All data needed to evaluate the conclusions in the paper are present in the paper and/or the Supplementary Materials. Additional data related to this paper may be requested from the lead contact, Jixi Li.

## Supplementary Materials

**Figure S1.**
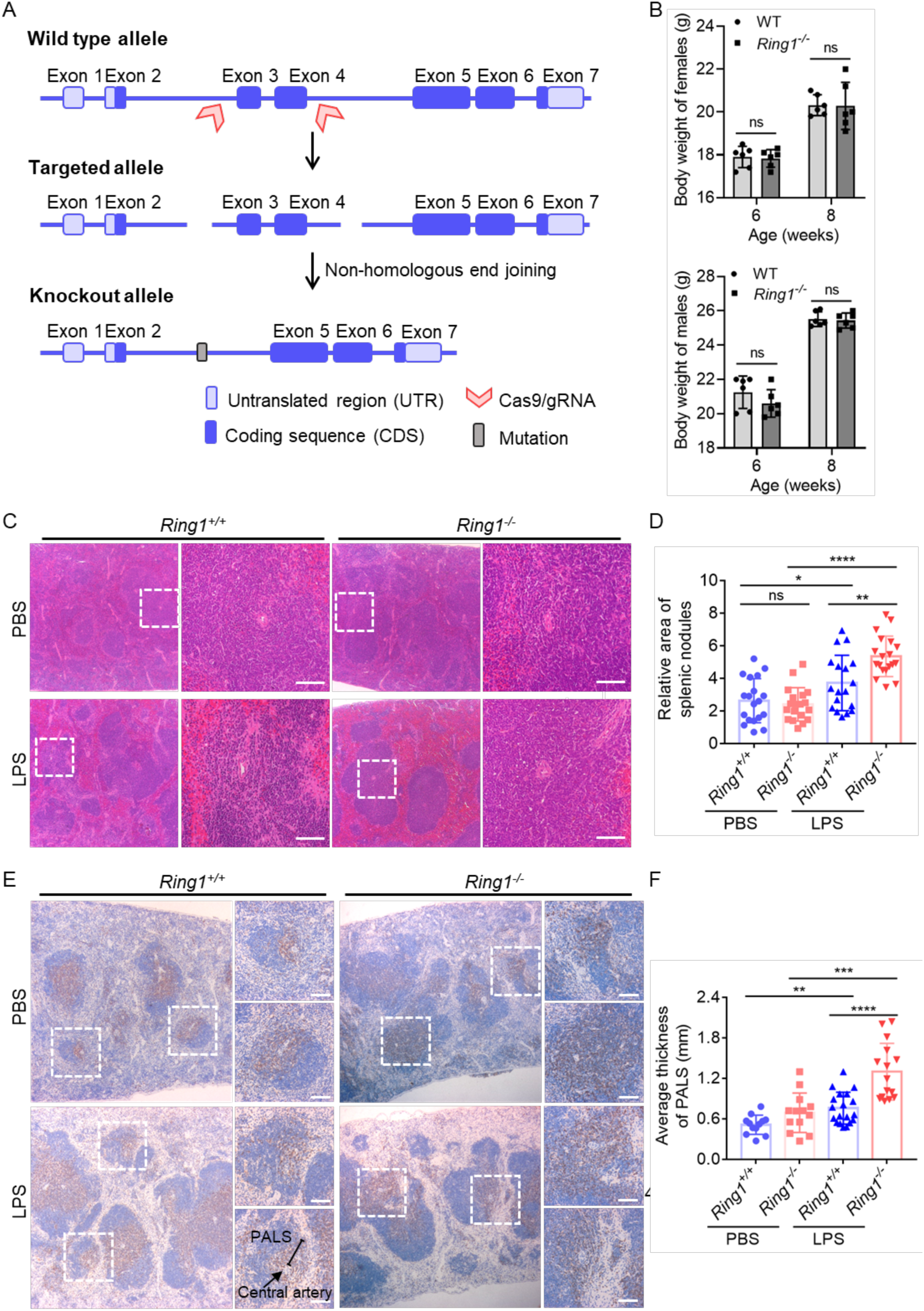
RING1 deficiency enhances LPS-induced inflammation. A. Schematic representation of the CRISPR gene-editing method in mice. B. Quantification analysis of mouse body weight. C. Histopathology of spleen sections from WT or *Ring1^-/-^* mice treated as in Figure 2A were examined by H&E staining. Scale bar, 400 μm. D. Quantification of relative areas of splenic nodules in panel C. E. The protein expression of CD4 in spleen tissues was analyzed by immunohistochemistry (IHC). Scale bar, 400 μm. F. Quantification of the periarterial lymphatic sheaths (PALS), as indicated by the white box in panel E.

**Figure S2.**
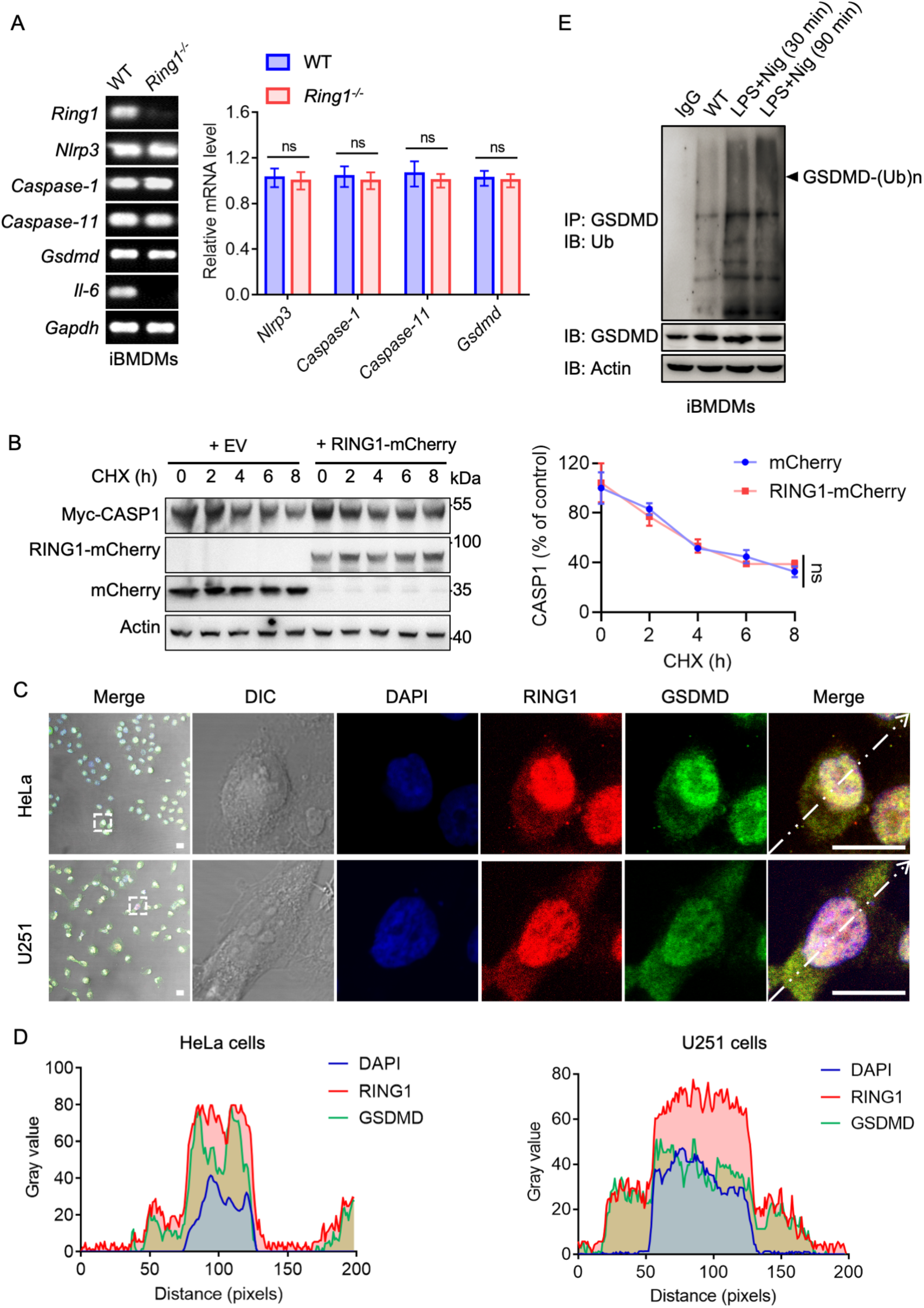
Endogenous RING1 colocalizes with GSDMD. A. mRNA levels of each gene in WT and *Ring1^-/-^* iBMDMs. The quantitative analysis was performed in the right panel. B. Detection of Myc-CASP1 half-life incubated with 30 μM CHX for indicated times. Quantification analysis was performed in the right panel. C. HeLa cells and U251 cells were fixed, and protein localization was visualized using specific antibodies. Nuclei were stained with Hoechst 33342 (Beyotime) for 10 min at room temperature (RT). Scale bar, 10 μm. D. Fluorescence intensity was plotted along the arrows in panel C. E. iBMDMs were primed with LPS for 4 h followed by Nig treatment or no treatment for 2 h. Cells were subjected to GSDMD pull-down and IB.

**Figure S3.**
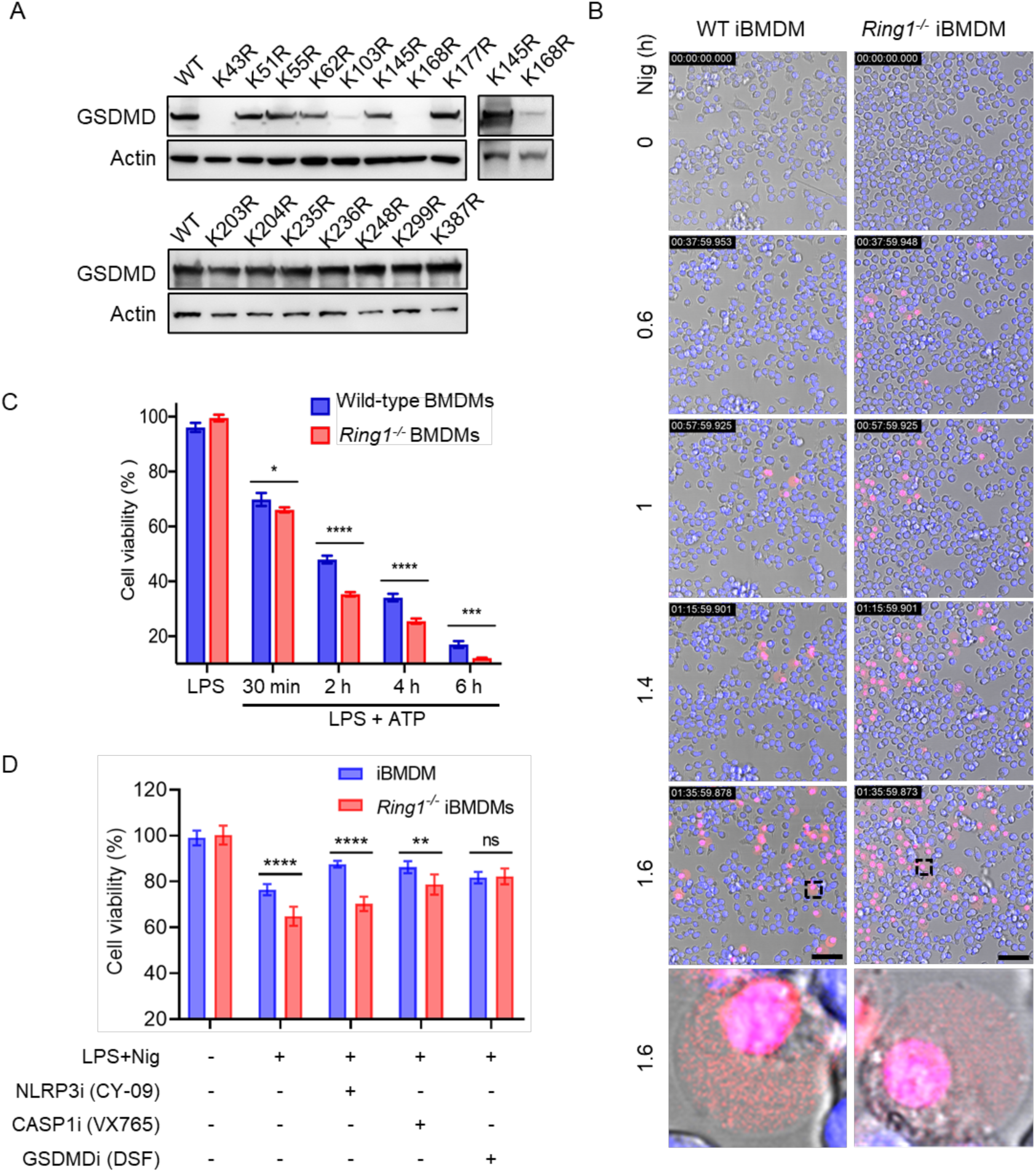
*Ring1* knockout increased pyroptosis in macrophages. A. HEK293T cells were transfected with HA-GSDMD or mutants and the proteins were detected by IB. B. WT or *Ring1^-/-^* iBMDMs were cultured in treated with 20 μM Nig for the indicated times. Hoechst 33342 live-cell nuclear dye and 5 μg/mL PI were added into the culture medium. Scale bar, 50 μm. C. The cell viability rates of BMDMs were treated as in Figure 5C. D. WT and *Ring1^-/-^* iBMDMs were pre-treated with 1 μg/mL LPS and the indicated inhibitors (CY-09 10 μM, VX765 20 μM, DSF 40μM) for 2 hours, followed by treatment of 20 μM Nig for 2 hours. Cell viability rates was determined by CCK-8 assay. Data represent three independent experiments and error bars represent the standard deviation (SD). Statistical analysis was performed using Two-way ANOVA (C-D). *, *p* < 0.05; **, *p* < 0.01; ***, *p* < 0.001; ****, *p* < 0.0001; ns, not significant.

**Figure S4.**
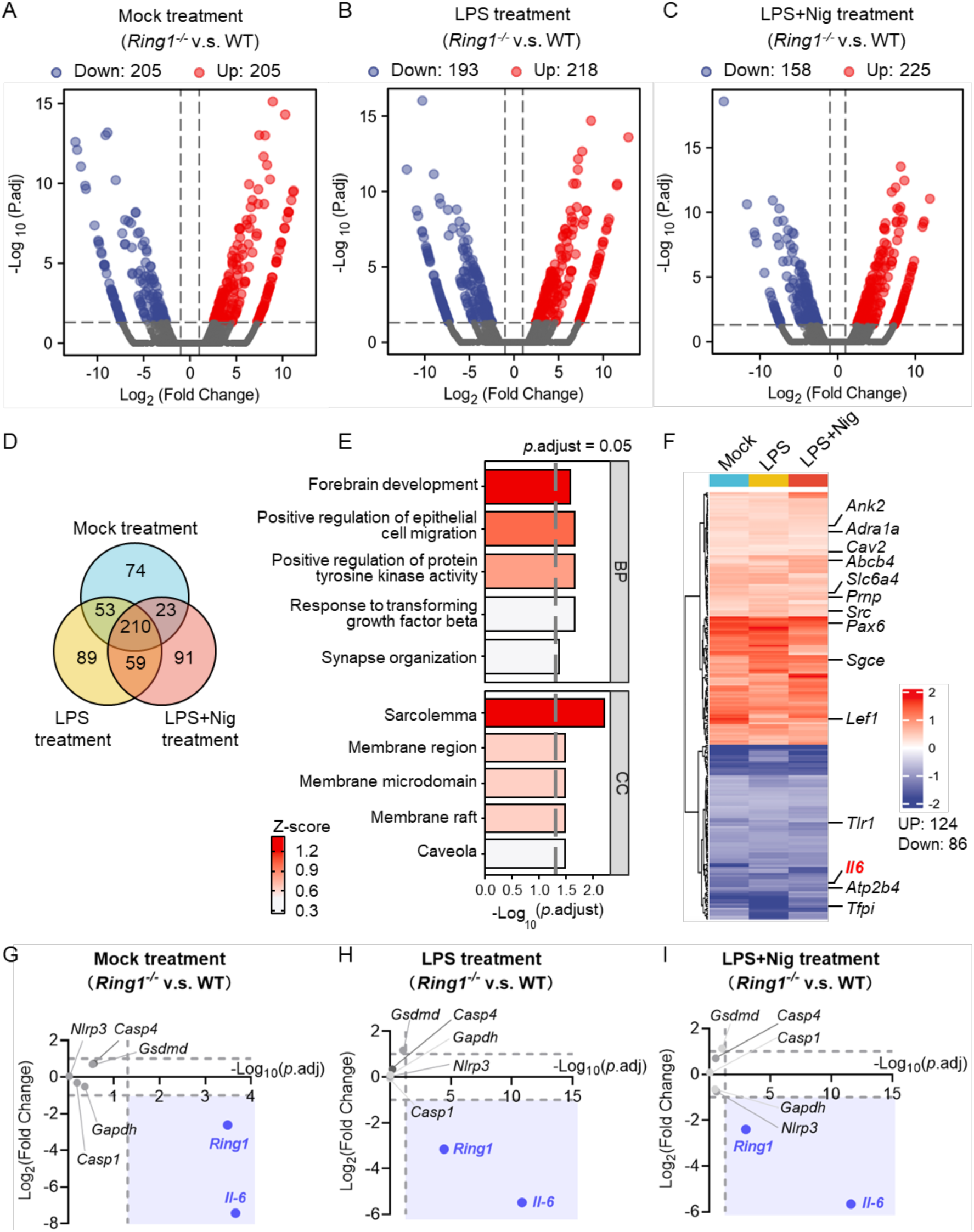
RNA-seq analysis of *Ring1^-/-^* and WT macrophages upon different treatments. A-C. Volcano plot indicating DEGs between *Ring1^-/-^* and WT iBMDMs under different treatments. D. Venn diagrams showing the most DEGs shared between the groups of different treatments. The overlapping sections represent the count of shared genes, while the distinct sections indicate the count of unique genes. E. GO category distribution of the 210 DEGs in the panel D. CC, cellular component. BP, biological process. F. Heatmap of up-and down-regulated DEGs shared across different treatments. The genes highlighted in black on the heatmap represent those enriched in the GO terms. G-I. mRNA expression levels of pyroptosis-related genes in iBMDMs under different conditions.

## References

Balakrishnan, M., and Kenworthy, A.K. (2024). Lipid Peroxidation Drives Liquid-Liquid Phase Separation and Disrupts Raft Protein Partitioning in Biological Membranes. J Am Chem Soc 146, 1374–1387. 10.1021/jacs.3c10132.

Beckwith, K.S., Beckwith, M.S., Ullmann, S., Saetra, R.S., Kim, H., Marstad, A., Asberg, S.E., Strand, T.A., Haug, M., Niederweis, M., et al. (2020). Plasma membrane damage causes NLRP3 activation and pyroptosis during Mycobacterium tuberculosis infection. Nat Commun 11, 2270. 10.1038/s41467-020-16143-6.

Blackledge, N.P., and Klose, R.J. (2021). The molecular principles of gene regulation by Polycomb repressive complexes. Nat Rev Mol Cell Biol 22, 815–833. 10.1038/s41580-021-00398-y.

Briukhovetska, D., Dorr, J., Endres, S., Libby, P., Dinarello, C.A., and Kobold, S. (2021). Interleukins in cancer: from biology to therapy. Nat Rev Cancer 21, 481–499. 10.1038/s41568-021-00363-z.

Chai, Q., Wang, L., Liu, C.H., and Ge, B. (2020). New insights into the evasion of host innate immunity by Mycobacterium tuberculosis. Cell Mol Immunol 17, 901–913. 10.1038/s41423-020-0502-z.

Chai, Q., Yu, S., Zhong, Y., Lu, Z., Qiu, C., Yu, Y., Zhang, X., Zhang, Y., Lei, Z., Qiang, L., et al. (2022). A bacterial phospholipid phosphatase inhibits host pyroptosis by hijacking ubiquitin. Science 378, eabq0132. 10.1126/science.abq0132.

Chandra, P., Grigsby, S.J., and Philips, J.A. (2022). Immune evasion and provocation by Mycobacterium tuberculosis. Nat Rev Microbiol 20, 750–766. 10.1038/s41579-022-00763-4.

Cohen, S.B., Gern, B.H., Delahaye, J.L., Adams, K.N., Plumlee, C.R., Winkler, J.K., Sherman, D.R., Gerner, M.Y., and Urdahl, K.B. (2018). Alveolar Macrophages Provide an Early Mycobacterium tuberculosis Niche and Initiate Dissemination. Cell Host Microbe 24, 439–446 e434. 10.1016/j.chom.2018.08.001.

del Mar Lorente, M., Marcos-Gutierrez, C., Perez, C., Schoorlemmer, J., Ramirez, A., Magin, T., and Vidal, M. (2000). Loss- and gain-of-function mutations show a polycomb group function for Ring1A in mice. Development 127, 5093–5100. 10.1242/dev.127.23.5093.

Devant, P., and Kagan, J.C. (2023). Molecular mechanisms of gasdermin D pore-forming activity. Nat Immunol 24, 1064–1075. 10.1038/s41590-023-01526-w.

Eto, H., Kishi, Y., Yakushiji-Kaminatsui, N., Sugishita, H., Utsunomiya, S., Koseki, H., and Gotoh, Y. (2020). The Polycomb group protein Ring1 regulates dorsoventral patterning of the mouse telencephalon. Nat Commun 11, 5709. 10.1038/s41467-020-19556-5.

Gao, W., Li, Y., Liu, X., Wang, S., Mei, P., Chen, Z., Liu, K., Li, S., Xu, X.W., Gan, J., et al. (2022). TRIM21 regulates pyroptotic cell death by promoting Gasdermin D oligomerization. Cell Death Differ 29, 439–450. 10.1038/s41418-021-00867-z.

He, W.T., Wan, H., Hu, L., Chen, P., Wang, X., Huang, Z., Yang, Z.H., Zhong, C.Q., and Han, J. (2015). Gasdermin D is an executor of pyroptosis and required for interleukin-1beta secretion. Cell Res 25, 1285–1298. 10.1038/cr.2015.139.

Hong, S.G., Ashby, J.W., Kennelly, J.P., Wu, M., Steel, M., Chattopadhyay, E., Foreman, R., Tontonoz, P., Tarling, E.J., Turowski, P., et al. (2024). Mechanosensitive membrane domains regulate calcium entry in arterial endothelial cells to protect against inflammation. J Clin Invest 134. 10.1172/JCI175057.

Hu, J.J., Liu, X., Xia, S., Zhang, Z., Zhang, Y., Zhao, J., Ruan, J., Luo, X., Lou, X., Bai, Y., et al. (2020). FDA-approved disulfiram inhibits pyroptosis by blocking gasdermin D pore formation. Nat Immunol 21, 736–745. 10.1038/s41590-020-0669-6.

Huang, Y., Xu, W., and Zhou, R. (2021). NLRP3 inflammasome activation and cell death. Cell Mol Immunol 18, 2114–2127. 10.1038/s41423-021-00740-6.

Kayagaki, N., Stowe, I.B., Lee, B.L., O’Rourke, K., Anderson, K., Warming, S., Cuellar, T., Haley, B., Roose-Girma, M., Phung, Q.T., et al. (2015). Caspase-11 cleaves gasdermin D for non-canonical inflammasome signalling. Nature 526, 666–671. 10.1038/nature15541.

Kuang, S., Zheng, J., Yang, H., Li, S., Duan, S., Shen, Y., Ji, C., Gan, J., Xu, X.W., and Li, J. (2017). Structure insight of GSDMD reveals the basis of GSDMD autoinhibition in cell pyroptosis. Proc Natl Acad Sci U S A 114, 10642–10647. 10.1073/pnas.1708194114.

Liu, S., Jiang, M., Wang, W., Liu, W., Song, X., Ma, Z., Zhang, S., Liu, L., Liu, Y., and Cao, X. (2018). Nuclear RNF2 inhibits interferon function by promoting K33-linked STAT1 disassociation from DNA. Nat Immunol 19, 41–52. 10.1038/s41590-017-0003-0.

Liu, Y.C., Penninger, J., and Karin, M. (2005). Immunity by ubiquitylation: a reversible process of modification. Nat Rev Immunol 5, 941–952. 10.1038/nri1731.

Luchetti, G., Roncaioli, J.L., Chavez, R.A., Schubert, A.F., Kofoed, E.M., Reja, R., Cheung, T.K., Liang, Y., Webster, J.D., Lehoux, I., et al. (2021). Shigella ubiquitin ligase IpaH7.8 targets gasdermin D for degradation to prevent pyroptosis and enable infection. Cell Host Microbe 29, 1521–1530 e1510. 10.1016/j.chom.2021.08.010.

Mei, P., Xie, F., Pan, J., Wang, S., Gao, W., Ge, R., Gao, B., Gao, S., Chen, X., Wang, Y., et al. (2021). E3 ligase TRIM25 ubiquitinates RIP3 to inhibit TNF induced cell necrosis. Cell Death Differ 28, 2888–2899. 10.1038/s41418-021-00790-3.

Mooney, E.C., and Sahingur, S.E. (2021). The Ubiquitin System and A20: Implications in Health and Disease. J Dent Res 100, 10–20. 10.1177/0022034520949486.

Nowarski, R., Jackson, R., and Flavell, R.A. (2017). The Stromal Intervention: Regulation of Immunity and Inflammation at the Epithelial-Mesenchymal Barrier. Cell 168, 362–375. 10.1016/j.cell.2016.11.040.

Peeters, J.G.C., and DuPage, M. (2021). A PRC1-RNF2 knockout punch for cancer. Nat Cancer 2, 996–997. 10.1038/s43018-021-00270-0.

Pierce, S.B., Stewart, M.D., Gulsuner, S., Walsh, T., Dhall, A., McClellan, J.M., Klevit, R.E., and King, M.C. (2018). De novo mutation in RING1 with epigenetic effects on neurodevelopment. Proc Natl Acad Sci U S A 115, 1558–1563. 10.1073/pnas.1721290115.

Ryan, C.W., Regan, S.L., Mills, E.F., McGrath, B.T., Gong, E., Lai, Y.T., Sheingold, J.B., Patel, K., Horowitz, T., Moccia, A., et al. (2024). RING1 missense variants reveal sensitivity of DNA damage repair to H2A monoubiquitination dosage during neurogenesis. Nat Commun 15, 7931. 10.1038/s41467-024-52292-8.

Shapouri-Moghaddam, A., Mohammadian, S., Vazini, H., Taghadosi, M., Esmaeili, S.A., Mardani, F., Seifi, B., Mohammadi, A., Afshari, J.T., and Sahebkar, A. (2018). Macrophage plasticity, polarization, and function in health and disease. J Cell Physiol 233, 6425–6440. 10.1002/jcp.26429.

Shen, J., Li, P., Shao, X., Yang, Y., Liu, X., Feng, M., Yu, Q., Hu, R., and Wang, Z. (2018). The E3 Ligase RING1 Targets p53 for Degradation and Promotes Cancer Cell Proliferation and Survival. Cancer Res 78, 359–371. 10.1158/0008-5472.CAN-17-1805.

Shi, J., Zhao, Y., Wang, K., Shi, X., Wang, Y., Huang, H., Zhuang, Y., Cai, T., Wang, F., and Shao, F. (2015). Cleavage of GSDMD by inflammatory caspases determines pyroptotic cell death. Nature 526, 660–665. 10.1038/nature15514.

Shi, Y., Yang, Y., Xu, W., Shi, D., Xu, W., Fu, X., Lv, Q., Xia, J., and Shi, F. (2022). E3 ubiquitin ligase SYVN1 is a key positive regulator for GSDMD-mediated pyroptosis. Cell Death Dis 13, 106. 10.1038/s41419-022-04553-x.

Shima, H., Takamatsu-Ichihara, E., Shino, M., Yamagata, K., Katsumoto, T., Aikawa, Y., Fujita, S., Koseki, H., and Kitabayashi, I. (2018). Ring1A and Ring1B inhibit expression of Glis2 to maintain murine MOZ-TIF2 AML stem cells. Blood 131, 1833–1845. 10.1182/blood-2017-05-787226.

Taru, V., Szabo, G., Mehal, W., and Reiberger, T. (2024). Inflammasomes in chronic liver disease: Hepatic injury, fibrosis progression and systemic inflammation. J Hepatol. 10.1016/j.jhep.2024.06.016.

van der Poll, T., Shankar-Hari, M., and Wiersinga, W.J. (2021). The immunology of sepsis. Immunity 54, 2450–2464. 10.1016/j.immuni.2021.10.012.

Voncken, J.W., Roelen, B.A., Roefs, M., de Vries, S., Verhoeven, E., Marino, S., Deschamps, J., and van Lohuizen, M. (2003). Rnf2 (Ring1b) deficiency causes gastrulation arrest and cell cycle inhibition. Proc Natl Acad Sci U S A 100, 2468–2473. 10.1073/pnas.0434312100.

Wang, Y., Li, Q., Zhang, J., Liu, P., Zheng, H., Chen, L., Wang, Z., Tan, C., Zhang, M., Zhang, H., et al. (2023). Ring1a protects against colitis through regulating mucosal immune system and colonic microbial ecology. Gut Microbes 15, 2251646. 10.1080/19490976.2023.2251646.

Wu, Y., Zhang, J., Yu, S., Li, Y., Zhu, J., Zhang, K., and Zhang, R. (2022). Cell pyroptosis in health and inflammatory diseases. Cell Death Discov 8, 191. 10.1038/s41420-022-00998-3.

Xia, S., Zhang, Z., Magupalli, V.G., Pablo, J.L., Dong, Y., Vora, S.M., Wang, L., Fu, T.M., Jacobson, M.P., Greka, A., et al. (2021). Gasdermin D pore structure reveals preferential release of mature interleukin-1. Nature 593, 607–611. 10.1038/s41586-021-03478-3.

Zhang, Z., Luo, L., Xing, C., Chen, Y., Xu, P., Li, M., Zeng, L., Li, C., Ghosh, S., Della Manna, D., et al. (2021). RNF2 ablation reprograms the tumor-immune microenvironment and stimulates durable NK and CD4(+) T-cell-dependent antitumor immunity. Nat Cancer 2, 1018–1038. 10.1038/s43018-021-00263-z.

Zhu, K., Li, J., Li, J., Sun, J., Guo, Y., Tian, H., Li, L., Zhang, C., Shi, M., Kong, G., and Li, Z. (2020). Ring1 promotes the transformation of hepatic progenitor cells into cancer stem cells through the Wnt/beta-catenin signaling pathway. J Cell Biochem 121, 3941–3951. 10.1002/jcb.29496.

